# NIK stabilization reprograms the pancreas toward chronic pancreatitis and pancreatic lipomatosis

**DOI:** 10.64898/2026.10.07.757290

**Authors:** Ziwei Du, Lilli Weber, Ulrike F.G. Büttner, Melanie Gerstenlauer, Uta Manfras, Katja Steiger, Thomas Metzler, Lap Kwan Chan, Medhanie Mulaw, Miltiadis Tsesmelis, Thomas Wirth

## Abstract

Pancreatic lipomatosis, characterized by progressive adipose replacement of the exocrine pancreas, is a known consequence of chronic pancreatitis. Despite its clinical relevance, the mechanisms driving lipomatosis remain poorly understood. Here, we used a pancreas-specific NF-κB-inducing kinase (NIK) stabilization mouse model to investigate how sustained non-canonical NF-κB signaling contributes to chronic pancreatitis and pancreatic lipomatosis. NIK stabilization initiated a rapid and fully penetrant pancreatic remodeling program, leading to acinar cell death already by the end of the first week of age. Deep learning on high-definition spatial transcriptomics revealed a progressive loss of acinar identity resulting in acinar-to-ductal metaplasia (ADM) formation and identified GPA33 as a novel surface marker of ADMs. Over time, ADM lesions underwent apoptosis, leading to progressive loss of pancreatic parenchyma, organ atrophy, and extensive adipose replacement. Mechanistically, *Relb* deletion fully rescued the NIK-induced pancreatic phenotype, demonstrating that chronic pancreatitis and lipomatosis require RelB-dependent non-canonical NF-κB signaling in this model. Together, these findings define a RelB-dependent pathway linking sustained NIK stabilization to chronic pancreatitis and pancreatic lipomatosis. Our study provides a model for studying the development of pancreatic lipomatosis and for exploring potential therapeutic approaches.

## Introduction

Chronic pancreatitis is a progressive, irreversible inflammatory disease of the pancreas that leads to persistent pain, loss of exocrine function, and a significant reduction in quality of life [1]. Importantly, chronic pancreatitis is a strong risk factor for pancreatic ductal adenocarcinoma (PDAC), with studies indicating an increased, up to 22-fold, chance of developing PDAC compared to the general population [2]. Thus, it is of utmost importance that the underlying mechanisms are thoroughly understood to aid the development of more effective treatments.

Pancreatic lipomatosis, also referred to as fatty replacement, fatty infiltration, or pancreatic steatosis, describes the accumulation of fat within the pancreatic parenchyma, often replacing exocrine tissue [3]. Chronic pancreatitis is recognized as one of the predisposing factors for pancreatic lipomatosis, with the process typically involving atrophy of acinar tissue followed by fat infiltration or replacement. The relationship between chronic pancreatitis and subsequent lipomatosis is complex, with evidence suggesting that chronic ductal obstruction and acinar cell loss can lead to progressive fat replacement of the exocrine pancreas [4].

The NF-κB family of transcription factors is central to the regulation of inflammation and immune responses, with two main activation routes, the canonical and the non-canonical pathways [5]. The non-canonical NF-κB pathway is regulated by NF-κB-inducing kinase (NIK), which is normally kept at low levels through continuous degradation by the TRAF-cIAP complex. Upon stimulation from upstream effectors, the TRAF complex is degraded, allowing NIK to accumulate and phosphorylate the downstream kinase IKKα. Activated IKKα then results in proteolytic processing of p100 into p52, allowing the RelB:p52 dimer to translocate to the nucleus and regulate gene expression [5].

The canonical NF-κB pathway has been extensively studied in the context of inflammation and pancreatic diseases, including pancreatitis and pancreatic cancer. However, the role of the non-canonical NF-κB pathway in these conditions is less well understood. It has been previously suggested that loss of acinar cell IKKα can trigger spontaneous and progressive pancreatitis in mice, although this was attributed to defective autophagy rather than the non-canonical pathway [6]. In other organs, such as the liver, non-canonical NF-κB signaling has been implicated in fat accumulation and metabolic regulation, but direct links to pancreatic fat infiltration remain to be fully established [7,8].

To investigate the role of NIK and non-canonical NF-κB signaling in the pancreas, the NIK^ΔT3^ mouse model was used. In this mouse line, the TRAF3-binding domain of NIK is deleted, resulting in stabilization and constitutive activation of the non-canonical pathway [9]. This persistent activation led to the rapid onset of acinar-to-ductal metaplasia (ADM) as early as two days after birth, progressing to severe pancreatitis with nearly complete replacement of the exocrine pancreas by ADMs by three weeks of age. By 20 weeks, ADM structures underwent cell death, and the pancreas was entirely replaced by fatty tissue, a process known as pancreatic lipomatosis, which was completely dependent on the non-canonical NF-κB pathway. To our knowledge, this is the first demonstration that constitutive activation of the non-canonical NF-κB pathway can fully recapitulate the features of chronic pancreatitis and pancreatic lipomatosis with 100% penetrance, highlighting a direct and potent role for this signaling axis in pancreatic disease.

## Results

### Pancreas-specific NIK stabilization triggers early ADM formation and fibrotic reaction

To investigate the function of NIK in the pancreas, we used a transgenic mouse model in which the TRAF3-binding domain of NIK was deleted, preventing its degradation by the TRAF complex (*R26-LSL-NIK^ΔT3^-IRES-eGFP*). The construct also drives eGFP expression to enable visualization of recombination events. We crossed these mice with *Pdx1-Cre* to generate *Pdx1-Cre; R26-LSL-NIK^ΔT3^-IRES-eGFP* (N^ΔT^C) animals, which led to constitutive NIK presence in the pancreas [9,10].

Mice were analyzed at three stages of pancreatic remodeling: the early acinar–ductal transition stage (2 days and 1 week), the intermediate remodeling stage (3 and 5 weeks), and the fat replacement stage (7, 10, and 20 weeks) (**Figure S1A**). Protein immunoblotting confirmed NIK expression in N^ΔT^C pancreata, whereas wild-type (WT) samples lacked detectable protein due to constitutive degradation (**Figure S1B**). In addition, GFP staining of 2-day-old pancreata showed a strong signal in N^ΔT^C acinar (amylase⁺) cells but not in WT controls, confirming efficient Cre-mediated recombination and NIK stabilization in the acinar compartment (**Figure S1C**).

Having validated the model, we next examined pancreatic morphology at the first stage. At postnatal day 2, N^ΔT^C pancreata appeared largely normal, with only rare abnormal duct-like structures (2%) detectable histologically (**Figure 1A, S1D**). However, by one week of age, these lesions had expanded significantly, encompassing nearly half of the pancreatic tissue (**Figure 1A**). Co-staining for amylase and CK19 revealed that these structures were double positive, confirming their acinar-to-ductal phenotype and identifying them as acinar-to-ductal metaplasia (ADM) lesions (**Figure 1B**).

**Figure 1:**
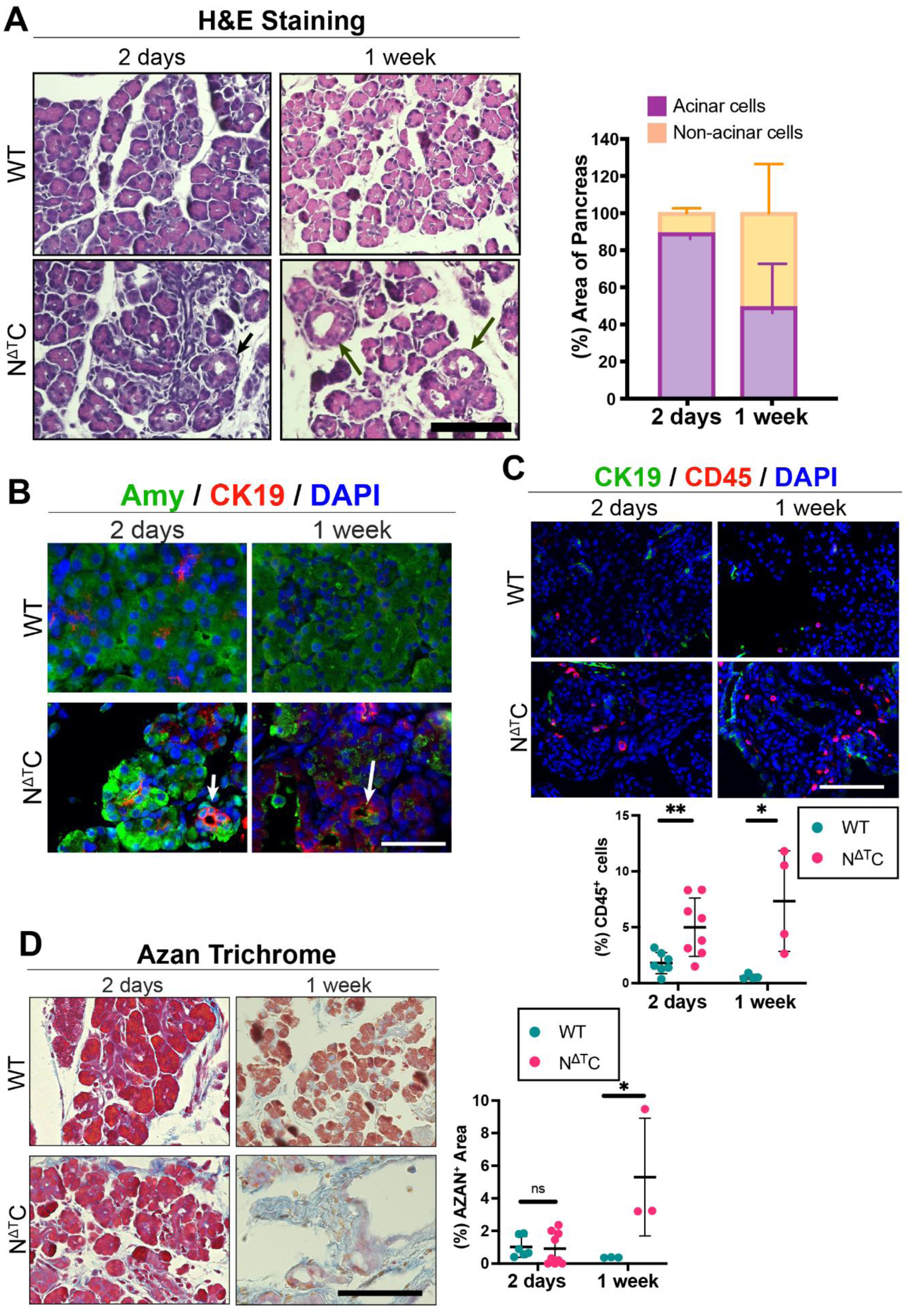
NIK activation promotes early pancreatic tissue remodeling, immune-cell infiltration, and fibrosis. **(A) Left:** Representative hematoxylin and eosin (H&E) staining of pancreatic sections from WT and N^ΔT^C mice at 2 days and 1 week. **Right:** Quantification of acinar and non-acinar areas as percentages of the total pancreatic tissue area at the indicated time points. N≥4 mice/group. P<0.05, Welch’s t-test. **(B)** Representative immunofluorescence staining of pancreatic sections from WT and N^ΔT^C mice at 2 days and 1 week for amylase (Amy, green), CK19 (red), and DAPI (blue). Scale bar: 50 μm. **(C) Top:** Representative immunofluorescence staining of pancreatic sections from WT and N^ΔT^C mice at 2 days and 1 week for CK19 (green), CD45 (red), and DAPI (blue). **Bottom:** Quantification of CD45⁺ cells as a percentage of total cells. N≥4 mice/group. 2 days: Welch’s t-test; 1 week: Unpaired t-test. **(D) Left:** Representative Azan trichrome staining of pancreatic sections from WT and N^ΔT^C mice at 2 days and 1 week. **Right:** Quantification of fibrotic area as a percentage of the total pancreatic tissue area. N≥3 mice/group. 2 days: Unpaired t-test. 1 week: Welch’s t-test. Dot plots represent individual mice. Data are presented as mean ± SD. ns: p > 0.05, *p < 0.05, **p < 0.01. Scale bars in (A), (C), and (D): 100 μm.

To better understand the cellular dynamics driving this transition, we examined the proliferation rate of acinar and ADM cells. WT amylase⁺ cells showed a higher proliferative rate than N^ΔT^C acinar cells (**Figure S1E**). Although ADMs exhibit elevated proliferation compared to acinar cells, our analysis was performed at postnatal day 2, when the pancreas is undergoing rapid growth [11]. At this stage, N^ΔT^C acinar cells have already begun adopting ADM-like features, and this phenotypic shift is likely associated with a reduction in proliferation.

We next assessed the inflammatory and fibrotic response in the pancreas. CD45 immunostaining revealed a clear increase in immune cell infiltration in N^ΔT^C pancreata at both 2 days and 1 week of age compared to WT (**Figure 1C**). In parallel, we observed more αSMA⁺ area, indicative of activated fibroblasts, which was modest at 2 days but became pronounced by 1 week (**Figure S1F**). Consistent with these findings, AZAN trichrome staining demonstrated significantly greater collagen deposition in N^ΔT^C pancreata at 1 week (**Figure 1D**). Together, these results indicate that NIK activation induces early acinar-to-ductal metaplasia accompanied by immune cell infiltration, fibroblast activation, and extracellular matrix deposition, all being key histopathological features of chronic pancreatitis [12].

### Spatial transcriptomics and deep learning approaches identify apoptotic signature and surface markers of late ADMs

To gain mechanistic insight into the processes described above, we performed high-definition spatial transcriptomics (Visium HD) on pancreata from WT and N^ΔT^C mice at 1 week of age **(Figure 2A**). This approach integrates histological and transcriptional information, enabling cell-type annotation within tissue architecture. Using hematoxylin and eosin (H&E) images from Visium HD, we manually annotated spatial spots based on morphological features (**Figure S2A**).

**Figure 2:**
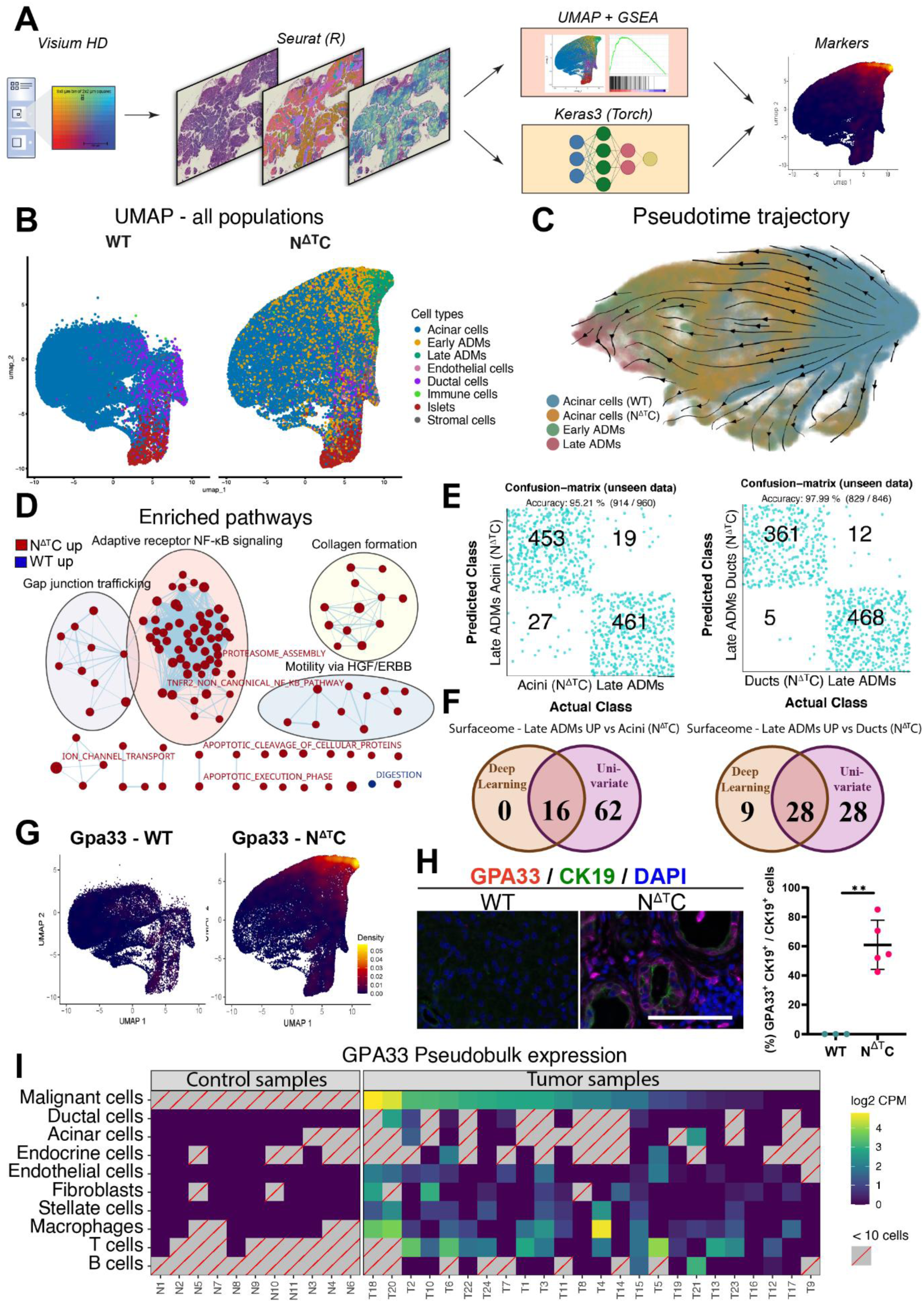
Spatial transcriptomics and deep learning characterize the acinar-to-ductal metaplasia trajectory in N^ΔT^C pancreata. **(A)** Schematic representation of the computational analysis followed. **(B)** UMAP of all annotated cell populations from WT and N^ΔT^C pancreata. **(C)** Pseudotime trajectory projected onto the UMAP of WT acinar cells, N^ΔT^C acinar cells, early ADMs, and late ADMs subset. Diffusion pseudotime values were derived using Slingshot and used as input for CellRank2 to infer the trajectory across the UMAP. **(D)** Deregulated pathways between WT and N^ΔT^C acinar cells. Gene set enrichment analysis (GSEA) was performed and the resulting pathways were clustered using Cytoscape. Red indicates pathways upregulated in N^ΔT^C acinar cells, and blue indicates pathways upregulated in WT acinar cells. **(E)** Confusion matrices from the deep learning classifiers for two comparisons: N^ΔT^C acinar cells versus late ADMs, and N^ΔT^C ductal cells versus late ADMs. For the comparisons involving acinar cells, the acinar population was randomly subsampled to match the cell number of late ADMs, and the analysis was repeated five times using five independent subsets. For these acinar cell comparisons, the mean of each classification is shown. **(F)** Venn diagrams showing the overlap between the top predictors from the deep learning models and the differentially expressed genes from the univariate analysis, for each of the two comparisons. Deep learning top predictors were extracted using integrated gradients, and differentially expressed genes were identified using the Wilcoxon rank-sum test. **(G)** Feature plot showing Gpa33 expression on the UMAP, split into WT and N^ΔT^C cells. **(H) Left**: immunofluorescence staining against GPA33 and CK19 in WT and N^ΔT^C pancreata. **Right**: quantification of the staining (N≥3, Welch’s t-test). Scale bars: 100 μm. **(I)** GPA33 pseudobulk expression (log2 CPM) across cell populations and patients in CRA001160, split into tumor-adjacent and tumor samples. Grey squares with a red diagonal mark populations that were absent or below the cell-number cutoff.

During the annotation, we noted heterogeneity among ADM lesions. Some structures showed only initial ductal conversion, with most acinar architecture preserved, whereas others exhibited complete restructuring into ductal-like formations. We therefore categorized these first lesions as early ADMs, representing transitional acinar regions, and the latter as late ADMs, representing fully established ductal-like structures (**Figure S2B**).

Following quality control and normalization, we generated a Uniform Manifold Approximation and Projection (UMAP) to visualize transcriptomic relationships across all annotated spots (**Figure 2B and S2C**). To examine whether acinar cells and ADMs form a continuous transcriptional trajectory, we subset acinar cells from WT and N^ΔT^C mice, as well as the early and late ADMs, constructed a diffusion map from their principal components, and computed diffusion pseudotime (**Figure 2C**). The four populations resolved along a single trajectory, ordered from acinar WT cells through N^ΔT^C acinar cells, early ADMs and, finally, late ADMs. This arrangement is consistent with a continuous transition from normal acinar cells toward fully established ductal-like structures.

To identify the early transcriptional alterations driving ADM initiation, we compared the transcriptomes of WT and N^ΔT^C acinar cells. Gene set enrichment analysis (GSEA) revealed activation of the non-canonical NF-κB pathway in N^ΔT^C acinar cells, consistent with NIK stabilization (**Figure 2D**). In addition, pathways related to collagen formation were enriched, corroborating our earlier histological evidence of fibroblast activation and collagen deposition. We also observed deregulation of ion channel transport activity, a phenomenon frequently associated with chronic pancreatitis [13]. In parallel, the digestive enzyme-related pathway was downregulated, reflecting the loss of acinar identity, characteristic of ADM progression. Finally, we detected enrichment of HGF/MET-driven motility signaling together with deregulated Gap junction trafficking. HGF and its receptor MET are induced in the pancreas during pancreatitis as part of the regenerative response [14,15], and both processes align with the loss of polarity and the reorganization of cell-cell contacts that accompany ADM [16].

Notably, N^ΔT^C acinar cells displayed an upregulation of pathways associated with apoptosis and cell death (**Figure 2D**). To validate this finding, we performed cleaved caspase-3 immunostaining and TUNEL assays, both of which confirmed an increased rate of apoptosis in N^ΔT^C pancreata compared to WT (**Figure S2D and S2E**). Consistent with these results, serum analysis from 2-day-old and 1-week-old N^ΔT^C mice revealed elevated levels of blood amylase, indicative of acinar cell injury (**Figure S2F**). These results suggest that NIK stabilization not only promotes ADM-associated transcriptional reprogramming but also triggers acinar cell death which together can drive the development of chronic pancreatitis.

To identify cell surface markers distinguishing late ADM lesions from other pancreatic populations, we developed a series of binary deep learning classifiers comparing late ADMs against WT acinar cells and WT ductal cells. To exclude the possibility that marker upregulation resulted from NIK stabilization itself rather than the ADM phenotype, we additionally included N^ΔT^C acinar and ductal cells as comparator groups. Each model demonstrated high predictive accuracy on independent-test data, with classification accuracy ranging from 95.2% to 99.6% (**Figure 2E and S2G**).

We next used the correlation coefficients from each trained predictor to extract surfaceome-associated genes linked to the late ADM class. These candidate markers were intersected with the top predictors identified by univariate differential expression analysis to obtain upregulated surfaceome features distinguishing late ADMs (**Figure 2F and S2H**). Intersecting all four pairwise comparisons yielded 8 surface markers reproducibly enriched in the late ADM population (**Figure S2I and S2J**). Among these markers, *Gpa33* emerged as the most robust marker, showing strong transcription in late ADM cells while remaining virtually absent in acinar and ductal populations (**Figure 2G**).

To validate GPA33 expression at the protein level, we performed immunofluorescence staining on pancreata from WT and N^ΔT^C mice. Consistent with the spatial transcriptomic data, GPA33 protein was detected on the surface of ADM cells in N^ΔT^C pancreata, whereas ductal cells in WT mice showed virtually no staining (**Figure 2H**).

To examine whether *GPA33* is also present in abnormal duct-like pancreatic cells in humans, we used the publicly available single-cell RNA-seq dataset from Peng et al., which includes tumor and control tissue from PDAC patients [17]. We compared malignant cells, that is, transformed duct-like cells potentially originating from ADMs, with acinar and ductal cells from both the tumor and control samples. *GPA33* expression was significantly enriched in the malignant cells compared to both acinar and ductal cells (**Figure 2I and S2K**). Consistent with the duct-like phenotype of these cells, *KRT19* showed the same enrichment pattern in malignant cells relative to acinar and ductal cells (**Figure S2L**). *KRT19* marks the pancreatic ductal lineage and is reactivated upon oncogenic KRAS signaling [18], supporting the interpretation that the malignant population reflects transformed duct-like cells.

Taken together, the high-definition spatial transcriptomic data revealed that consistent NIK signaling promotes acinar cell death and tissue remodeling, features associated with pancreatitis severity [19], and identified GPA33 as a surface marker of late ADMs.

### NIK stabilization drives ADM remodeling and a shift toward cell death over proliferation

We next analyzed the intermediate stage of pancreatic remodeling at 3 and 5 weeks of age. Histological examination at 3 weeks revealed nearly complete replacement of the exocrine pancreas by ADMs and development of chronic pancreatitis (**Figure 3A**). Similarly, pancreata at 5 weeks were histologically comparable, indicating persistence of this remodeling phase (**Figure 3A**).

**Figure 3:**
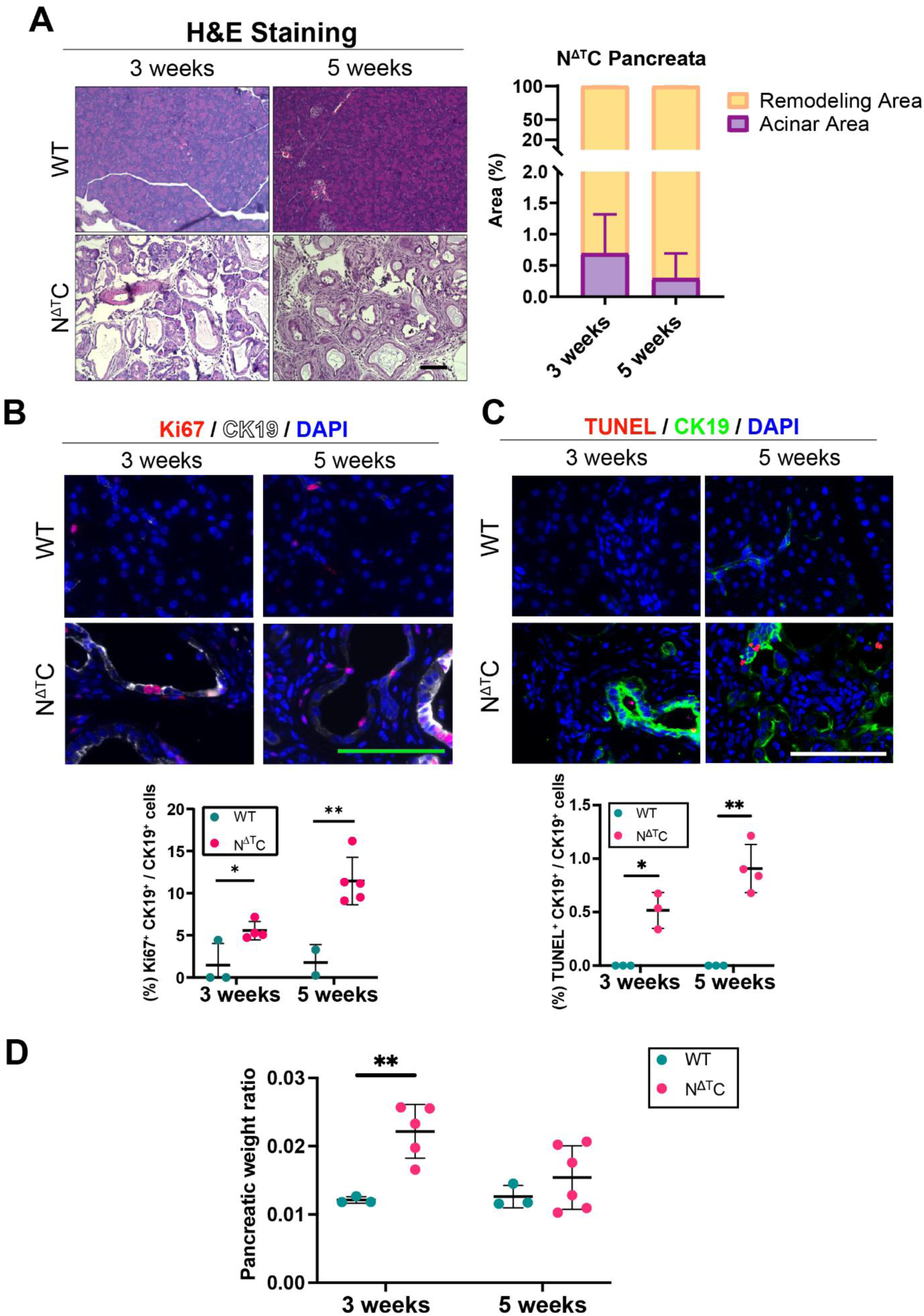
NIK activation drives chronic ADM remodeling and cell death. **(A) Left:** Representative hematoxylin and eosin (H&E) staining of pancreatic sections from WT and N^ΔT^C mice at 3 and 5 weeks. **Right:** Quantification of acinar and remodeling areas as percentages of the total pancreatic tissue area in N^ΔT^C pancreata at the indicated time points. N≥3 mice/group. **(B) Top:** Representative immunofluorescence staining of pancreatic sections from WT and N^ΔT^C mice at 3 and 5 weeks for Ki67 (red), CK19 (white), and DAPI (blue). **Bottom:** Quantification of Ki67⁺CK19⁺ cells as a percentage of total CK19⁺ cells. N≥2 mice/group. Unpaired t-test. **(C) Top:** Representative immunofluorescence staining of pancreatic sections from WT and N^ΔT^C mice at 3 and 5 weeks for TUNEL (red), CK19 (green), and DAPI (blue). **Bottom:** Quantification of TUNEL⁺CK19⁺ cells as a percentage of total CK19⁺ cells. N≥3 mice/group. Welch’s t-test. **(D)** Quantification of the pancreas-to-body weight ratio in WT and N^ΔT^C mice at 3 and 5 weeks. N≥3 mice/group. 3 weeks: Welch’s t-test; 5 weeks: Unpaired t-test. Dot plots represent individual mice. Data are presented as mean ± SD. *p < 0.05, **p < 0.01. Scale bars: 100 μm.

To characterize the stromal response, we performed α-SMA immunostaining and AZAN trichrome staining. Both analyses revealed extensive fibroblast activation and collagen deposition throughout the exocrine pancreas of N^ΔT^C mice, while WT pancreata remained negative (**Figure S3A and S3B**).

Examination of cell proliferation showed a significant increase in CK19⁺ cells in N^ΔT^C pancreata compared to WT (**Figure 3B**). In parallel, cleaved caspase 3 and TUNEL staining revealed apoptotic CK19⁺cells in N^ΔT^C pancreata, whereas WT ductal cells were negative (**Figure 3C and S3C**), indicating that both proliferative and apoptotic processes are active within the N^ΔT^C ductal cells.

To evaluate which process predominates at the organ level, we compared the pancreas-to-body weight ratios of WT and N^ΔT^C mice at these time points. At 3 weeks, N^ΔT^C pancreata were significantly enlarged relative to WT, consistent with the effect of active fibrosis. However, by 5 weeks, this ratio had declined in N^ΔT^C mice and no longer differed from WT, suggesting that acinar cell loss/atrophy progressively outweighed proliferation (**Figure 3D**).

### NIK stabilization leads to extensive adipose replacement and pancreatic atrophy

We next analyzed the late stage of pancreatic remodeling at 7, 10, and 20 weeks of age. Histological analysis at 7 weeks revealed that most of the pancreatic tissue remained affected by chronic pancreatitis. However, small islands of fat deposition were visible, covering approximately 3% of the total area (**Figure 4A**). Oil Red O staining confirmed the lipid nature of these cells, verifying that they represented adipocytes (**Figure 4B**). By 10 weeks, fat accumulation increased markedly, occupying about 20% of the pancreatic area, and by 20 weeks, adipocytes replaced nearly 70% of the tissue, indicating a progressive fatty replacement of the pancreas. At this late stage, adipocyte expansion was accompanied by reduction in the area covered by ADMs, fibrosis, and inflammation, suggesting that lipogenic remodeling gradually overtakes the chronic pancreatitis phenotype (**Figure 4C and S4A**).

**Figure 4:**
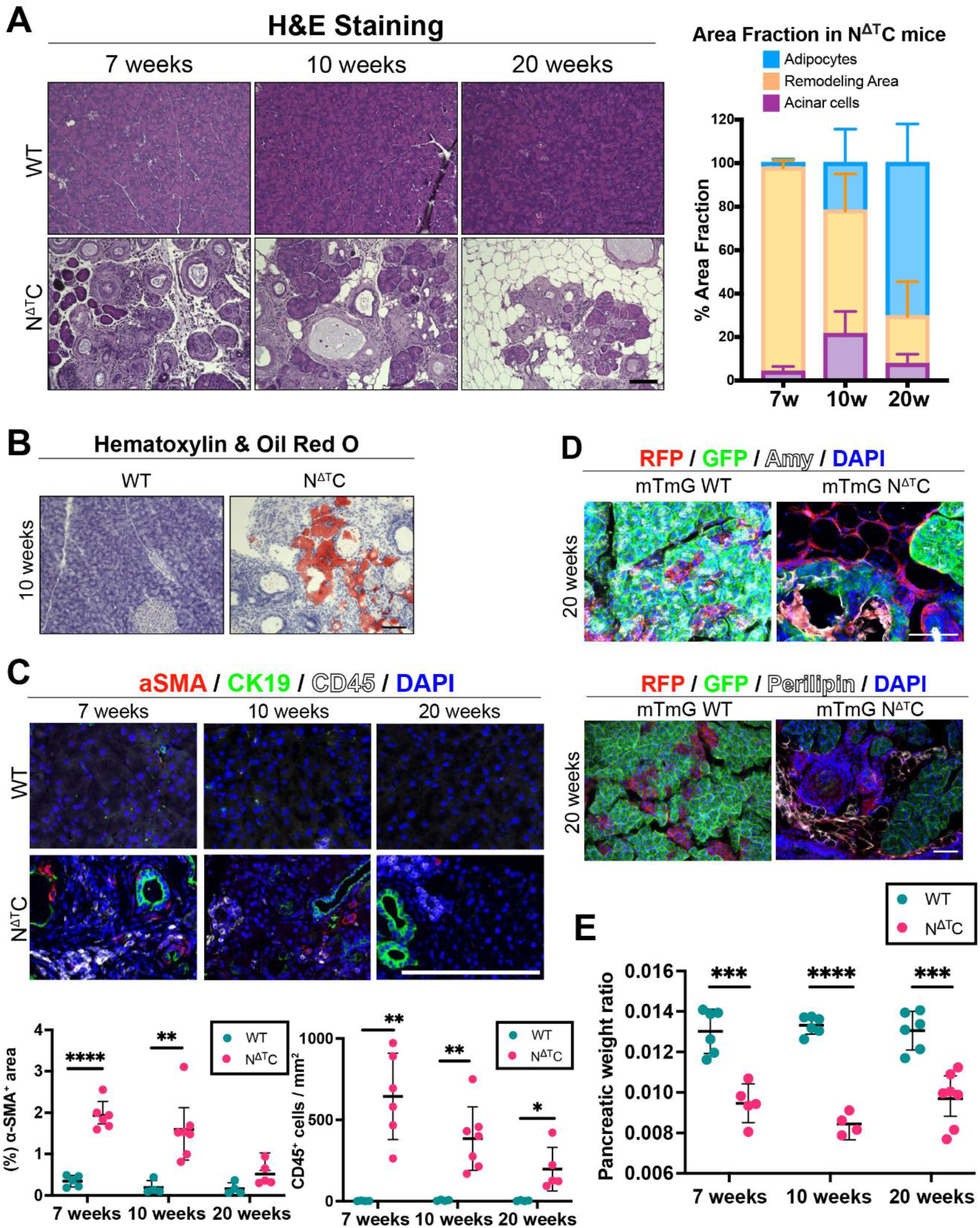
NIK activation leads to extensive adipose replacement and pancreatic atrophy. **(A) Left:** Representative hematoxylin and eosin (H&E) staining of pancreatic sections from WT and N^ΔT^C mice at 7, 10, and 20 weeks. **Right:** Quantification of acinar, adipose, and remodeling areas as percentages of the total pancreatic tissue area in N^ΔT^C mice at the indicated time points. N≥6 mice/group. **(B)** Representative hematoxylin and Oil Red O staining of pancreatic sections from WT and N^ΔT^C mice at 10 weeks, showing lipid accumulation in N^ΔT^C pancreata. **(C) Top:** Representative immunofluorescence staining of pancreatic sections from WT and N^ΔT^C mice at 7, 10, and 20 weeks for α-SMA (red), CK19 (green), CD45 (white), and DAPI (blue). **Bottom Left:** Quantification of α-SMA-positive area as a percentage of the total tissue area. N≥6 mice/group. 7, 20 weeks: Unpaired t-test; 10 weeks: Mann-Whitney test. **Bottom Right:** Quantification of CD45⁺ immune cells per mm². N≥6 mice/group. 7, 10 weeks: Welch’s t-test; 20 weeks: Mann-Whitney test. **(D)** Representative lineage-tracing immunofluorescence images of pancreatic sections from mTmG WT and mTmG N^ΔT^C mice at 20 weeks. **Top:** RFP (red), GFP (green), amylase (white), and DAPI (blue). **Bottom:** RFP (red), GFP (green), perilipin (white), and DAPI (blue), showing the relationship between recombined pancreatic epithelial cells and adipocyte-containing regions. **(E)** Quantification of the pancreas-to-body weight ratio in WT and NΔTC mice at 7, 10, and 20 weeks. N≥6 mice/group. Unpaired t-test. Dot plots represent individual mice. Data are presented as mean ± SD. *p < 0.05, **p < 0.01, ***p < 0.001, ****p < 0.0001. Scale bars: 100 μm.

To determine the origin of the adipocytes, we used mTmG reporter mice under the control of the *Pdx1-Cre* promoter, where cells of pancreatic origin express EGFP and non-pancreatic cells express tdTomato (*ROSAmT/mG*). Perilipin staining of N^ΔT^C pancreata at 20 weeks revealed that the adipocytes were tdTomato^+^, indicating they were not derived from pancreatic lineages. These results argue against transdifferentiation of acinar or metaplastic cells and indicate a non-epithelial source for the fatty replacement (**Figure 4D**).

We next assessed the proliferative and apoptotic activity of ADM (CK19^+^) cells during this late stage (**Figure S4B, S4C and S4D**). At 7 weeks, ADM cells exhibited their highest proliferative activity, which progressively declined at 10 and 20 weeks. A similar pattern was observed for apoptosis, with elevated cell death at 7 weeks that decreased at later time points. Importantly, all analyses showed statistically significant differences between WT and N^ΔT^C pancreata at each stage.

Consistent with the progressive apoptosis, the pancreas-to-body weight ratio was reduced in N^ΔT^C mice starting from 7 weeks and remained lower than in WT controls (**Figure 4E**). This reduction likely reflects both the ongoing death and atrophy of ADM cells and the gradual replacement of pancreatic tissue with adipocytes, which contribute less to overall weight.

Chronic pancreatitis and the associated fatty replacement have been linked, in certain cases, to the development of type 3c diabetes, although this outcome is not universally observed [3,20]. To evaluate whether our model exhibited such metabolic alterations, we measured blood glucose levels, total body weight in WT and N^ΔT^C mice throughout their lifespan, serum insulin concentrations as well as synaptophysin^+^, insulin^+^ and glucagon^+^ covered area per pancreas. No significant differences were detected between the two groups, indicating that NIK stabilization-induced pancreatic remodeling does not impair glucose homeostasis (**Figure S4E-S4H**).

Collectively, our results show that sustained NIK stabilization results in near-complete replacement of pancreatic parenchyma by adipose tissue, modeling a distinct subtype of chronic pancreatitis with extensive fatty remodeling.

### NIK-driven chronic pancreatitis requires activation of the non-canonical NF-κB pathway

We showed above that constitutive NIK stabilization activates the non-canonical NF-κB pathway and its downstream effectors. However, as NIK is a serine/threonine kinase, it could potentially influence additional targets beyond this pathway. To determine whether activation of the non-canonical NF-κB pathway mediates this phenotype, we generated N^ΔT^RC mice (*Pdx1-Cre; R26-LSL-NIK^ΔT3^-IRES-eGFP; Relb^fl/fl^*), in which RelB, the key downstream effector of this pathway, is deleted.

To verify efficient recombination and transgene expression, we first examined GFP expression in N^ΔT^RC pancreata. Robust GFP staining was observed throughout the pancreas, confirming successful activation of the NIK^ΔT3^ transgene despite RelB deletion (**Figure 5A**). We then assessed RelB expression in WT, N^ΔT^C, and N^ΔT^RC mice. As expected, N^ΔT^C pancreata displayed strong nuclear RelB staining, indicative of active non-canonical NF-κB signaling, whereas RelB signal was virtually absent in both WT and N^ΔT^RC pancreata (**Figure 5B**), confirming efficient RelB deletion in the latter.

**Figure 5:**
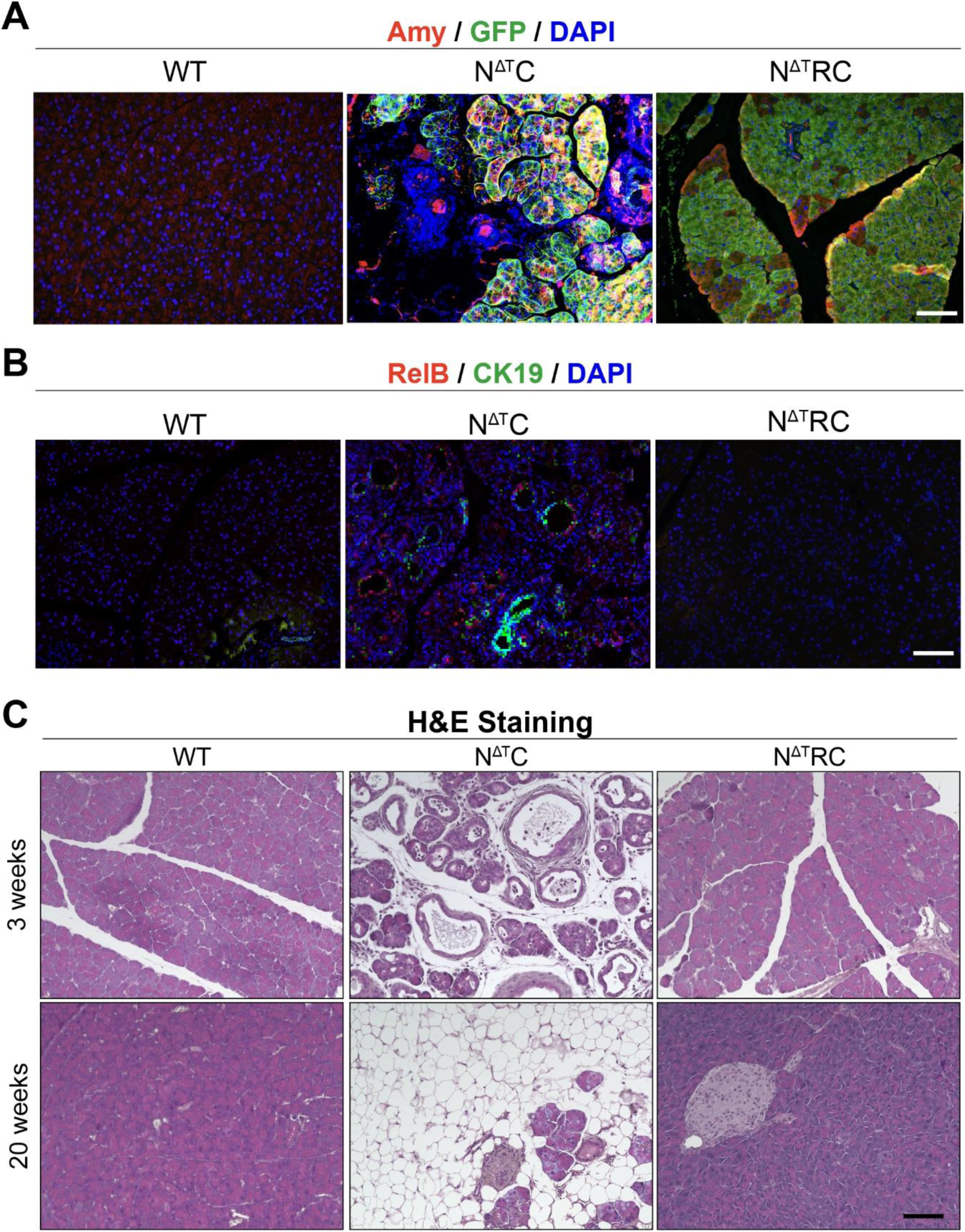
NIK-driven chronic pancreatitis requires activation of the non-canonical NF-κB pathway. **(A)** Representative immunofluorescence staining of pancreatic sections from WT, N^ΔT^C, and N^ΔT^RC mice for amylase (Amy, red), GFP (green), and DAPI (blue), showing preservation of acinar tissue in N^ΔT^RC mice. Scale bars: 100 μm. **(B)** Representative immunofluorescence staining of pancreatic sections from WT, N^ΔT^C, and N^ΔT^RC mice for RelB (red), CK19 (green), and DAPI (blue), confirming reduced RelB expression in N^ΔT^RC pancreata. Scale bars: 100 μm. **(C)** Representative hematoxylin and eosin (H&E) staining of pancreatic sections from WT, N^ΔT^C, and N^ΔT^RC mice at 3 and 20 weeks, showing restoration of pancreatic architecture following *Relb* deletion in the N^ΔT^C background. Scale bars: 100 μm.

Next, we compared the pancreatic phenotype across the different genotypes. By 3 weeks of age, N^ΔT^C mice exhibited severe chronic pancreatitis characterized by extensive ADM formation, increased proliferation, fibrosis, and immune cell infiltration. In contrast, N^ΔT^RC pancreata remained histologically indistinguishable from WT, with no evidence of ADM, fibrosis, or inflammation (**Figure 5C, S5A - S5C**). Consistent with these findings, the pancreas-to-body weight ratio was significantly increased in N^ΔT^C mice but remained at WT levels in N^ΔT^RC animals (**Figure S5D**).

Importantly, this protective effect persisted at later stages of disease progression. Histological assessment at 20 weeks demonstrated suppression of pancreatic abnormalities, fibrosis, inflammation or alterations to pancreas-to-body weight ratio in N^ΔT^RC mice (**Figure 5C, S5A - S5C**). Together, these findings demonstrate that RelB-dependent non-canonical NF-κB signaling is required for NIK-driven chronic pancreatitis and pancreatic lipomatosis, establishing this pathway as the mediator of the phenotype.

## Discussion

Chronic pancreatitis is a progressive inflammatory disorder associated with tissue remodeling and an increased risk of pancreatic cancer. One potential consequence of this process is pancreatic lipomatosis, a recognized but poorly understood condition in which pancreatic parenchyma is replaced by adipose tissue, and which is over-represented in pancreatic ductal adenocarcinoma, with reported odds of approximately sixfold [21]. Notably, hepatocyte-specific NIK overexpression triggers macrophage-dependent injury and fibrosis in the liver, while hepatic NIK activity is required for the development of hepatic steatosis [22,23]. In the pancreas, it has been shown that NIK deletion accelerates KRAS-driven tumorigenesis in association with remodeling of the tumor microenvironment, establishing that this kinase is functionally relevant in pancreatic tissue [24]. In the present study, we used a pancreas-specific NIK stabilization model to investigate the contribution of non-canonical NF-κB signaling to pancreatic remodeling. Our findings demonstrate that NIK stabilization drives a highly penetrant and progressive disease characterized by ADM formation, chronic pancreatitis, and pancreatic lipomatosis.

Already by the end of the first week of age (early transition stage), N^ΔT^C pancreata displayed ADM formation, increased apoptosis, and immune-cell infiltration, demonstrating that NIK stabilization is sufficient to initiate pancreatic remodeling shortly after birth. The formation of ADM lesions is consistent with the well-established property of acinar cells to undergo stress-induced reprogramming [16]. Correspondingly, transcriptomic analyses revealed deregulation of the digestive-enzyme and ion-transport pathways, indicating an early loss of acinar-cell identity and polarity [25]. Together with acinar cell apoptosis and inflammation, these findings support a model in which NIK-driven non-canonical NF-κB signaling promotes tissue remodeling. Finally, our findings are in line with previous studies linking NIK to inflammation and cell death in the liver [22].

GPA33, a transmembrane glycoprotein of the immunoglobulin superfamily, is normally expressed in blood and intestinal epithelial cells but not in pancreatic acinar cells [26,27]. Interestingly, our spatial transcriptomic analysis identified GPA33 as a surface marker of ADMs. In line with our results, it has recently been suggested that pancreatic metaplastic cells, after losing their acinar identity, partially switch on an intestinal differentiation program [28]. Importantly, our computational analysis of the human scRNA-seq PDAC study from Peng et al. showed that *GPA33* is also enriched in malignant cells compared to acinar and ductal cells, suggesting that its expression in deregulated pancreatic epithelium is conserved between mice and humans. Since GPA33 is expressed in 95% of colorectal cancers and is already being explored as a target in mice using CAR-T therapy, it may also be of interest as a marker or target in pancreatic diseases [29].

During the intermediate stage (3-5 weeks of age), N^ΔT^C pancreata were virtually replaced by ADMs and fibrosis. Although ADMs normally resolve after injury, this recovery did not occur in our mouse model, probably due to constant NIK stabilization or persistent inflammation. Interestingly, despite increased ADM cell proliferation, the reduction in pancreas mass suggests that apoptosis outpaced ADM expansion. We hypothesize that the continued loss of acinar and ADM cells sets the stage for later fatty replacement, consistent with observations from both mouse models and human studies [30,31].

During the late stage of disease progression (7–20 weeks of age), chronic pancreatitis transitioned into extensive pancreatic lipomatosis, resulting in near-complete replacement of the pancreatic parenchyma by adipose tissue. Although pancreatic lipomatosis has been described in patients with chronic pancreatitis, its occurrence is often heterogeneous and incompletely penetrant [3]. In contrast, sustained NIK stabilization drove a remarkably consistent progression toward end-stage fatty replacement in all animals examined, suggesting that activation of this axis is sufficient to orchestrate the full spectrum of disease.

A central question is the origin of the replacing adipocytes in pancreatic lipomatosis. In other models, pancreatic adipocytes can arise from the pancreatic epithelium, with acinar-to-adipocyte transdifferentiation reported after inactivation of *c-Myc* or *Gata6* [32,33]. In contrast, in our study, adipocytes in N^ΔT^C pancreata were uniformly tdTomato positive in our *ROSA26mT/mG* reporter mice, excluding a pancreatic epithelial origin. Our findings are consistent with pancreatic stellate cells and pancreas-resident mesenchymal stromal cells, which retain adipogenic potential and have been suggested as the main candidate source of ectopic pancreatic adipocytes [34]. This indicates that the fatty replacement arises from a non-epithelial source, potentially stromal or peripancreatic adipose expanding into parenchyma lost through acinar and ADM cell death, although the precise origin is yet to be defined.

With respect to endocrine compartment alterations, our analysis revealed a comparatively preserved islet architecture. In line with previous reports, constitutive NIK activation in β cells does not produce an overt diabetic phenotype, nor does β-cell-specific NIK deletion alter diabetes incidence [35,36]. Other studies, however, have reported endocrine involvement of NIK, with overexpression in β cells causing insulitis and spontaneous diabetes and α-cell-specific overexpression causing α-cell loss, reduced β-cell mass and pancreatitis [37,38]. These phenotypes are likely determined by the conditions a compartment is targeted and when the analysis is performed, indicating that the consequences of pathway deregulation are context-dependent.

Notably, non-canonical NF-κB pathway-associated mediators, such as IKKα and NIK, has been reported to exert biological functions beyond their established roles in non-canonical NF-κB signaling [6,8]. In the case of IKKα, its deletion led to chronic pancreatitis, associated with spontaneous acinar vacuolization, fibrosis and inflammation. The phenotype was attributed to p62 aggregate accumulation which was characterized as NF-κB-independent. Importantly, in our study, we examined whether our phenotype was NF-κB-dependent by performing genetic ablation of *Relb* in N^ΔΤ^C mice. Knockout of *Relb* completely rescued the development of pancreatitis, ADM formation, fibrosis, and pancreatic lipomatosis, demonstrating that the pathological effects of NIK in this model are mediated through the RelB-dependent non-canonical NF-κB pathway.

Taken together, the N^ΔT^C model recapitulates the full pathological sequence leading to pancreatic lipomatosis, from initial acinar injury and chronic pancreatitis, parenchymal loss and ultimately adipose replacement. Our study establishes a model for investigating the development of pancreatic lipomatosis and evaluating potential therapeutic approaches.

## Material and methods

### Mice

*Pdx1-Cre* mice were crossed with *R26-LSL-NIK^ΔT3^-IRES-eGFP* mice, in which deletion of the TRAF3-binding domain stabilizes NIK and constitutively activates non-canonical NF-κB signaling, to generate *Pdx1-Cre; R26-LSL-NIK^ΔT3^-IRES-eGFP* (N^ΔT^C) mice [9,10]. To examine the requirement for RelB in the phenotype, N^ΔT^C mice were crossed with *Relb^fl/fl^* mice to generate *Pdx1-Cre; R26-LSL-NIK^ΔT3^-IRES-eGFP; Relb^fl/fl^* (N^ΔT^RC) mice. For lineage-tracing experiments, ROSA^mT/mG^ reporter mice were included as indicated. Littermates lacking *Pdx1-Cre* served as wild-type (WT) controls. Both sexes were used. Mice were maintained on a C57BL/6 background under standard housing conditions at the animal facility of the University of Ulm and were analyzed at postnatal day 2, and weeks 1, 3, 5, 7, 10, or 20 of age, as indicated for each experiment. All animal experiments were performed in accordance with the ARRIVE guidelines, German animal-welfare legislation, and the approval of the responsible governmental authority (TVA1328).

### Tissue Processing and Histology

Pancreata were collected at the indicated time points, weighed, and normalized to body weight. Tissues were either snap-frozen for molecular analyses, fixed in 4% paraformaldehyde, fresh 10% neutral buffer formalin or embedded in optimal cutting temperature compound for cryosectioning. Formalin-fixed paraffin-embedded (FFPE) pancreatic sections (3 μm) were stained with hematoxylin and eosin (H&E) and Azan trichrome using established protocols. To assess fatty replacement, cryosections were stained with Oil Red O and counterstained with hematoxylin. Whole pancreatic H&E sections were used to quantify the relative area of normal exocrine tissue, ADM/remodeling, fibrosis, or adipose replacement, as appropriate. For field-based analyses, at least five randomly selected, non-overlapping fields per sample were quantified whenever tissue size permitted. Fewer fields were analyzed when the available tissue area was insufficient, as in pancreatic sections from mice aged 2 days or 1 week. Histopathological assessment was performed by veterinary pathologists using coded slides, with assessors blinded to genotype.

### Immunofluorescence (IF) and Immunohistochemistry (IHC)

FFPE sections were deparaffinized, subjected to heat-mediated antigen retrieval in citrate buffer (100 °C, 10 min). Sections were incubated with primary antibodies overnight at 4 °C. After washing with PBS, species-appropriate fluorescent secondary antibodies and DAPI were applied for 1 h at room temperature before mounting. For chromogenic IHC, endogenous peroxidase activity was quenched with 3% hydrogen peroxide after antigen retrieval. Sections were then blocked, incubated with primary antibodies, followed by biotinylated secondary antibodies and streptavidin; signals were visualized using AEC+ chromogen. Apoptotic cells were detected using the In Situ Cell Death Detection Kit (Roche #11684795910, Basel, Switzerland) according to the manufacturer’s instructions and, where indicated, co-stained with lineage markers. Antibodies and dilutions are listed in Supplementary Table S1. Images were acquired using identical exposure settings within each experiment and quantified using ImageJ/Fiji and QuPath.

### Western Blot

Protein was extracted from snap-frozen pancreatic tissue using SDS-based lysis buffer. Lysates were sonicated (40% amplitude; 3 × 7 s on/7 s off), resolved by SDS-PAGE, and transferred to membranes by wet or semi-dry transfer. Proteins were detected by enhanced chemiluminescence using a ChemiDoc MP Imaging System (Bio-Rad, Hercules, USA). Densitometric values were normalized to the indicated loading controls or to total protein levels. Antibodies and dilutions are listed in Supplementary Table S1.

### Sample Preparation and Visium HD Processing

Spatial transcriptomic profiling of FFPE pancreatic tissue sections was performed with the Visium HD Mouse Transcriptome, 6.5 mm kit (10x Genomics, PN-1000676) according to the manufacturer’s protocol. Tissue sections were processed using the Visium CytAssist instrument (10x Genomics, PN-1000441). Probe hybridization and ligation used Visium Mouse Transcriptome Probes v2, small (10x Genomics, PN-1000667).

### Library Construction and Sequencing

cDNA synthesis, amplification, and library construction were performed using the Visium HD Reagent Kit, Small (PN-1000668, 10x Genomics), with sample indexing performed using the Dual Index Kit TS Set A, 96 rxns (PN-1000251, 10x Genomics). Libraries were quantified using an Agilent Bioanalyzer and sequenced on an MGI DNBSEQ-G400 sequencing platform to a mean depth of approximately 1.42 billion read pairs per capture area.

### Data Processing and Alignment

Raw sequencing data were processed using the Space Ranger pipeline and the mm10 was used as reference genome. The resulting outs folder contained the filtered feature-barcode matrix, spatial images, and metadata necessary for downstream analysis that were used to generate the Seurat object.

### Spatial transcriptomics quality control and normalization

Quality control and normalization were performed on the spatial transcriptomics data prior to downstream analysis. Spots with fewer than 500 total RNA counts or with a mitochondrial RNA content exceeding 10% were excluded to remove low-quality observations. Gene expression values were normalized and variance-stabilized using the SCTransform method. Principal component analysis (PCA) was then conducted, and the first 25 principal components were used to generate a UMAP embedding for visualization and clustering.

### Cell Type Annotation

Cell annotation was performed using Loupe Browser (10x Genomics). Spots were classified based on their morphological characteristics, allowing identification of distinct cell populations. The separation between acinar cells, early acinar-to-ductal metaplasia (ADM), and late ADM regions is described in detail in the manuscript.

### Univariate analysis

To identify the top predictors distinguishing the two populations using a univariate model, we performed a differential expression analysis using the Wilcoxon rank-sum test. Features were ranked according to their adjusted p-values, and the top 5% of features with the lowest adjusted p-values were selected as the most significantly different between the populations. In addition to selecting the top features, we generated pre-ranked feature lists for each comparison. All detected features were ranked according to a composite score calculated as –10 × log₁₀(adjusted p-value), multiplied by the sign of the log₂ fold change (log₂FC). This approach assigns more extreme positive scores to significantly upregulated features and more extreme negative scores to significantly downregulated ones.

### Gene set enrichment analysis

Gene set enrichment analysis (GSEA) was performed using the pre-ranked feature lists generated for each comparison to identify significantly enriched or depleted gene sets. The analysis was conducted against the Reactome pathway database to determine biological processes and pathways associated with the observed expression differences. Enrichment results were visualized using Cytoscape software to facilitate network-based interpretation of pathway relationships.

### Trajectory and pseudotime analysis

Trajectory inference was performed on the subset of acinar cells from WT and N^ΔT^C mice together with the early and late ADMs. Starting from the Seurat object, we constructed a diffusion map from the first 35 principal components using *destiny* with a locally adaptive kernel width and density normalization, and retained the first 20 diffusion components as input for lineage inference. Lineages and pseudotime were then inferred with *Slingshot*, using acinar WT cells as the starting cluster. Cells were ordered along the resulting lineage by orthogonal projection onto the fitted principal curve.

To obtain a directional representation of this trajectory, the Slingshot pseudotime was used to derive a directed cell-cell transition matrix with *CellRank2*. A k-nearest-neighbour graph was computed on the principal component space (30 neighbours), and a PseudotimeKernel was applied with the Slingshot pseudotime as the ordering variable. The resulting transition matrix was projected onto the UMAP embedding to visualise the inferred direction of the acinar-to-ADM transition.

### Deep learning

We constructed deep learning models with an initial layer of all features (ranging from 11264 to 11671 nodes), a hidden layer of 512 nodes and a final output layer consisting of a single node for binary classification. Kernel weights were initialized using the He (Kaiming) initializer, and a Rectified Linear Unit (ReLU) activation function was applied to each layer. Model optimization was performed using a stochastic gradient descent method based on adaptive estimation (AdamW optimizer) with binary cross-entropy as the loss function. A weight decay of 1e-4 was applied to prevent overfitting. Additional L1 (Lasso) and L2 (Ridge) regularization were included, with coefficients of 1e-6 and 1e-4, respectively. For comparisons between acini (WT or N^ΔT^C) and late ADMs, we controlled for the marked imbalance in cell numbers by retaining all late ADM cells and repeatedly downsampling the acini population to a matched size (five independent subsamples with non-overlapping cells), reporting the features consistently identified across runs (intersection). Each model was trained using 60% of cells from both populations, with 20% reserved for validation. After each epoch, the model’s performance was evaluated on the validation set, and validation loss was recorded. If validation loss did not improve by at least 1e-4 for four consecutive epochs, the learning rate was reduced by half. If no improvement was observed for 12 epochs, training reverted to the best-performing epoch and was terminated.

### Deep learning top predictors

To estimate feature importance (correlation coefficients) from the deep learning models, we applied the Integrated Gradients (IG) method. This approach assigns an importance coefficient to each feature based on both its contribution magnitude and direction toward class prediction. IG computations were performed using 64 interpolation steps between the baseline and the actual input values, with a batch size of 32 for each analysis. The IG analysis was conducted separately for each pair of populations being compared. The top 5% of features with the highest absolute correlation coefficients were identified as the most influential features for class discrimination.

### Analysis of human PDAC single-cell RNA-seq data

Human pancreatic ductal adenocarcinoma (PDAC) single-cell RNA-seq data were obtained from the Genome Sequence Archive under accession CRA001160 [17], comprising 24 tumor samples and 11 tumor-adjacent tissues. The cell type annotation used was provided by the original authors. Ductal cell type 2 was relabeled as malignant and ductal cell type 1 as ductal, in line with the description in the original publication.

Pseudobulk profiles were generated by summing raw counts across all cells of a given cell type within each sample, using AggregateExpression in Seurat. Profiles derived from fewer than 10 cells were excluded from all downstream analyses. Differential expression was assessed with DESeq2. Malignant cells were compared against two reference populations, ductal and acinar cells, in three sample sets (all samples, tumor-adjacent samples, and tumor samples only), giving six comparisons in total. P values were adjusted for multiple testing using the Benjamini-Hochberg procedure, and an adjusted p value below 0.05 was considered significant. For visualization, expression was expressed as log2 counts per million calculated from the same pseudobulk matrix.

### Statistics

Statistical analyses were performed using GraphPad Prism unless otherwise stated. Data are presented as mean ± SD unless indicated otherwise. Each dot in the graphs represents one biological replicate, typically one mouse or one independently processed sample. The exact number of biological replicates is indicated in the corresponding figure legends. For comparisons between two groups, data distribution and variance were assessed before statistical testing. When data were normally distributed with comparable variance, an unpaired two-tailed Student’s t-test was used. When variance was unequal, Welch’s t-test was applied. For non-normally distributed data, the Mann–Whitney U test was used. For comparisons involving more than two groups with one independent variable, one-way ANOVA followed by Tukey’s multiple-comparison test was used when variances were homogeneous. If variances were unequal, Welch’s ANOVA was applied. All statistical tests were two-sided unless otherwise stated. Statistical significance was defined as follows: ns, P > 0.05; *P < 0.05; **P < 0.01; ***P < 0.001; ****P < 0.0001.

## Acknowledgments

The authors thank Prof. Dr. med Achim Weber for providing the facility and assistance on the immunohistochemical staining.

## Author contributions

Conceptualization, M.T., T.W., and Z.D.; methodology, M.T., Z.D. and U.B.; software, Z.D. and M.T.; validation, Z.D.; formal analysis, Z.D., M.T., T.M., K.S., and L.W.; investigation, Z.D., M.T., L.W., U.B., M.G., L.C., and U.M.; resources, T.W.; data curation, Z.D. and M.T.; writing – original draft, M.T. and Z.D.; Writing – review & editing, T.W. and M.T.; visualization, Z.D. and M.T.; supervision, T.W. and M.T.; project administration, Z.D., T.W. and M.T.; funding acquisition, T.W. All authors have read and agreed to the published version of the manuscript.

## Ethics approval

The study was performed according to ARRIVE guidelines and the German animal welfare legislation and approved by the responsible government agency under licence TVA1328.

## Data availability statement

The RNA-seq data supporting the findings of this study have been deposited in the GEO repository under accession GSE349857. The dataset includes both raw FASTQ files and processed data. Data will be made public upon manuscript publication or upon request. All other relevant data are available from the corresponding author upon reasonable request.

## Code availability

The custom code used for the deep learning classification, the Integrated Gradients gene ranking and the comparison with Seurat FindMarkers is publicly available on GitHub (https://github.com/miltos91/seurat-deep-learning-ig-findmarkers) under the MIT license, and the version used in this study is archived at Zenodo (https://doi.org/10.5281/zenodo.23210768). All other analyses were performed with published software packages as described in the Methods. Additional analysis.

