## Supplementary File for "NIK stabilization reprograms the pancreas toward chronic pancreatitis and pancreatic lipomatosis"

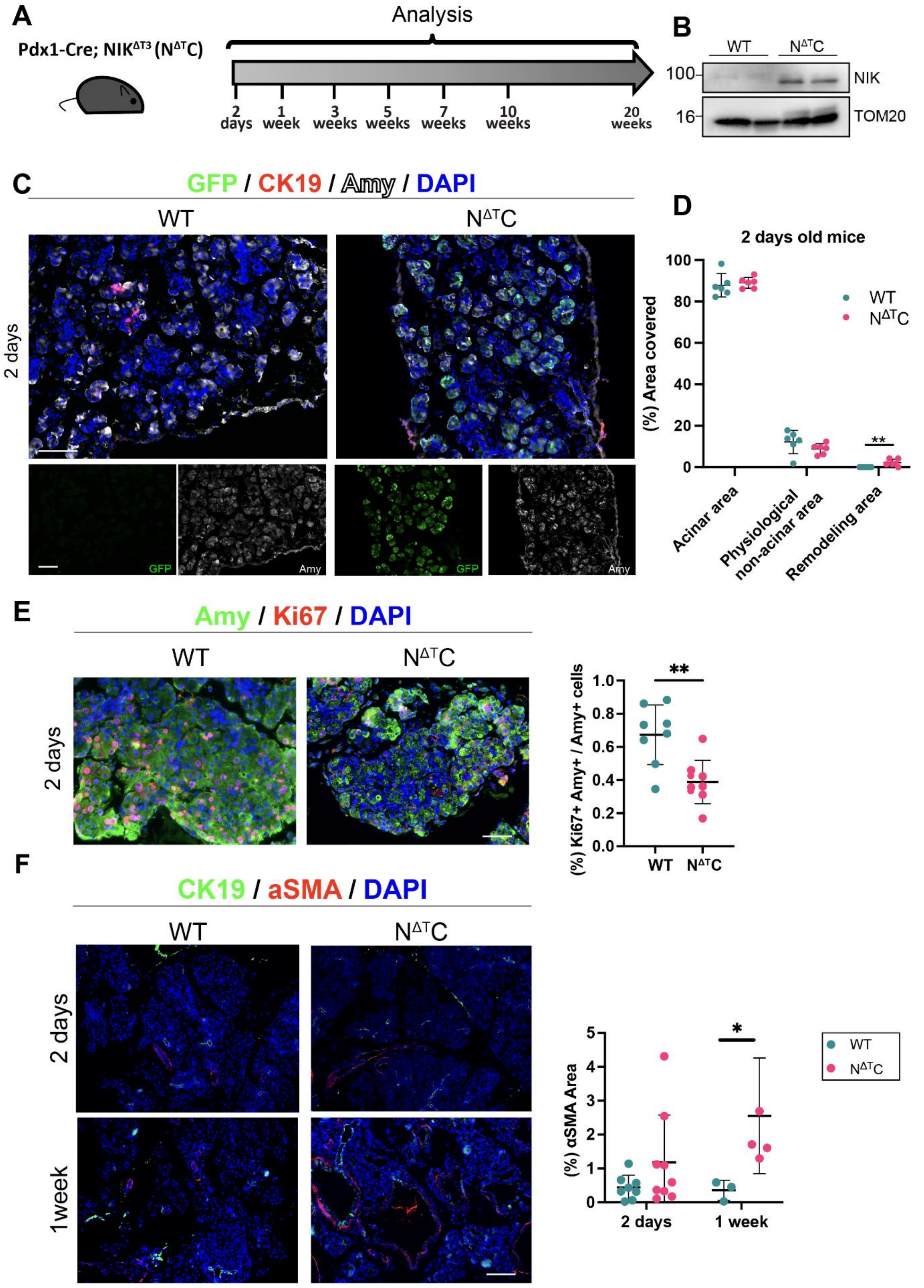


**Supplementary Figure S1. Experimental strategy and early pancreatic alterations following constitutive NIK activation.**

**(A)** Schematic representation of the experimental design. WT and Pdx1-Cre; NIK$^{\Delta T}$3 (N$^{\Delta T}$C) mice were analyzed at 2 days and at 1, 3, 5, 7, 10, and 20 weeks of age. **(B)** Western blot analysis of NIK protein levels in pancreatic tissue from WT and N$^{\Delta T}$C mice. TOM20 was used as a loading control. **(C)** Representative immunofluorescence staining of pancreatic sections from 2-day-old WT and N$^{\Delta T}$C mice for GFP (green), CK19 (red), amylase (Amy, white), and DAPI (blue). Lower panels show the GFP and amylase channels separately. **(D)** Quantification of acinar, non-acinar, and remodeling areas as percentages of the total pancreatic tissue area in 2-day-old WT and N$^{\Delta T}$C mice. N ≥ 8 mice/group. Unpaired t-test. **(E) Left:** Representative immunofluorescence staining of pancreatic sections from 2-day-old WT and N$^{\Delta T}$C mice for amylase (Amy, green), Ki67 (red), and DAPI (blue). **Right:** Quantification of Ki67$^+$Amy$^+$DAPI$^+$ cells as a percentage of total Amy$^+$DAPI$^+$ cells. N ≥ 8 mice/group. Unpaired t-test. **(F) Left:** Representative immunofluorescence staining of pancreatic sections from WT and NΔTC mice at 2 days and 1 week for CK19 (green), α-SMA (red), and DAPI (blue). **Right:** Quantification of α-SMA-positive area as a percentage of the total pancreatic tissue area. N ≥ 4 mice/group. Mann-Whitney test.

Dot plots represent individual mice. Data are presented as mean ± SD. *p < 0.05, **p < 0.01. Scale bars: 100 μm.

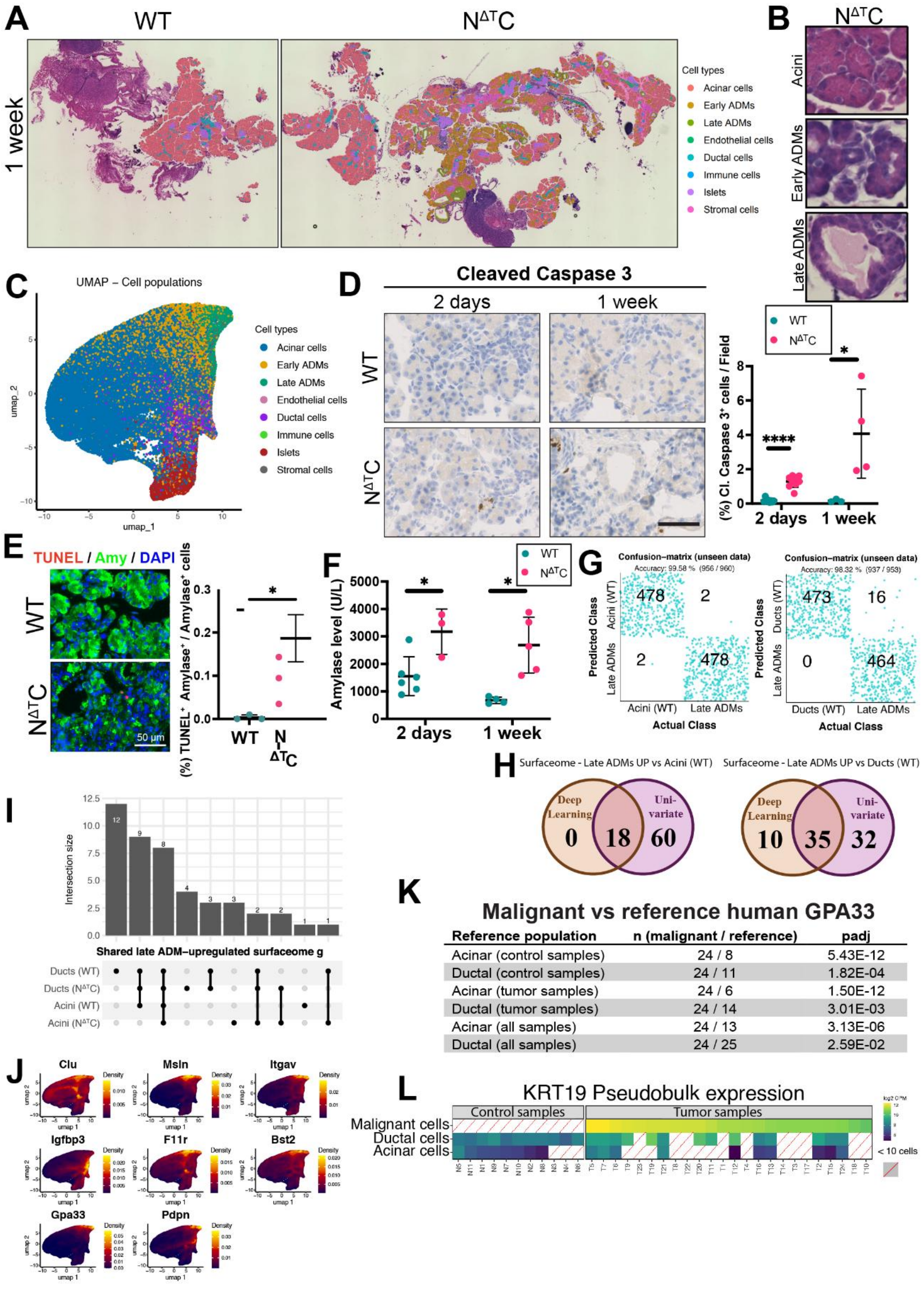


**Supplementary Figure S2**. **Spatial transcriptomic characterization following constitutive NIK activation.**

**(A)** H&E staining from wild-type and N$^{\Delta T}$C pancreata used for the spatial transcriptomics. The cell annotations are overlayed on cell by color. **(B)** Representative images of the annotated cell populations in $N^{\Delta T}C$ pancreata, showing acinar cells, early ADMs, and late ADMs. **(C)** UMAP of all annotated cell populations from wild-type and $N^{\Delta T}C$ pancreata combined. **(D) Left**: Cleaved caspase 3 staining of WT and $N^{\Delta T}C$ pancreata at 2 days and 1 week of age. Scale bars: 100 µm. **Right**: Quantification of the staining. N≥4 mice/group. Welch's t-test. **(E) Left**: immunofluorescence staining against TUNEL and α-amylase in WT and $N^{\Delta T}C$ pancreata at 1 week of age. Scale bars: 50 µm. **Right**: quantification of the staining N≥3 mice/group. Welch's t-test. **(F)** Amylase levels in the blood of WT and $N^{\Delta T}C$ mice at 2 days and 1 week of age. N≥4 mice/group. Unpaired t-test. **(G)** Confusion matrices from the deep learning classifiers for two comparisons: WT acinar cells versus late ADMs, wild-type ductal cells versus late ADMs. For the comparisons involving acinar cells, the acinar population was randomly subsampled to match the cell number of late ADMs, and the analysis was repeated five times using five independent subsets. For these comparisons, the mean of each classification is shown. **(H)** Venn diagrams showing the overlap between the top predictors from the deep learning models and the differentially expressed genes from the univariate analysis, for each of the two comparisons. Deep learning top predictors were extracted using integrated gradients, and differentially expressed genes were identified using the Wilcoxon rank-sum test. **(I)** UpSet plot showing the intersections of the shared late ADM-upregulated surfaceome genes across the four comparisons (Ducts WT, Ducts $N^{\Delta T}C$, Acini WT, Acini $N^{\Delta T}C$) **(J)** Feature plots showing the expression of all eight detected markers on the UMAP. **(K)** Differential expression of *GPA33* between malignant cells and reference epithelial populations in human PDAC (CRA001160). Six comparisons are shown, using ductal and acinar cells from all samples, from tumor-adjacent tissues, and from tumour samples only. Log2 fold changes with Benjamini-Hochberg adjusted p values from DESeq2 on patient-level pseudobulk profiles, with group sizes given as the number of pseudobulk samples. **(L)** *KRT19* pseudobulk expression (log2 CPM) across cell populations and patients in CRA001160, split into tumor-adjacent and tumor samples. Grey squares with a red diagonal mark populations that were absent or below the cell-number cutoff.

Dot plots represent individual mice. Data are presented as mean ± SD. *p < 0.05, ****p < 0.0001. Scale bars: 100 µm.

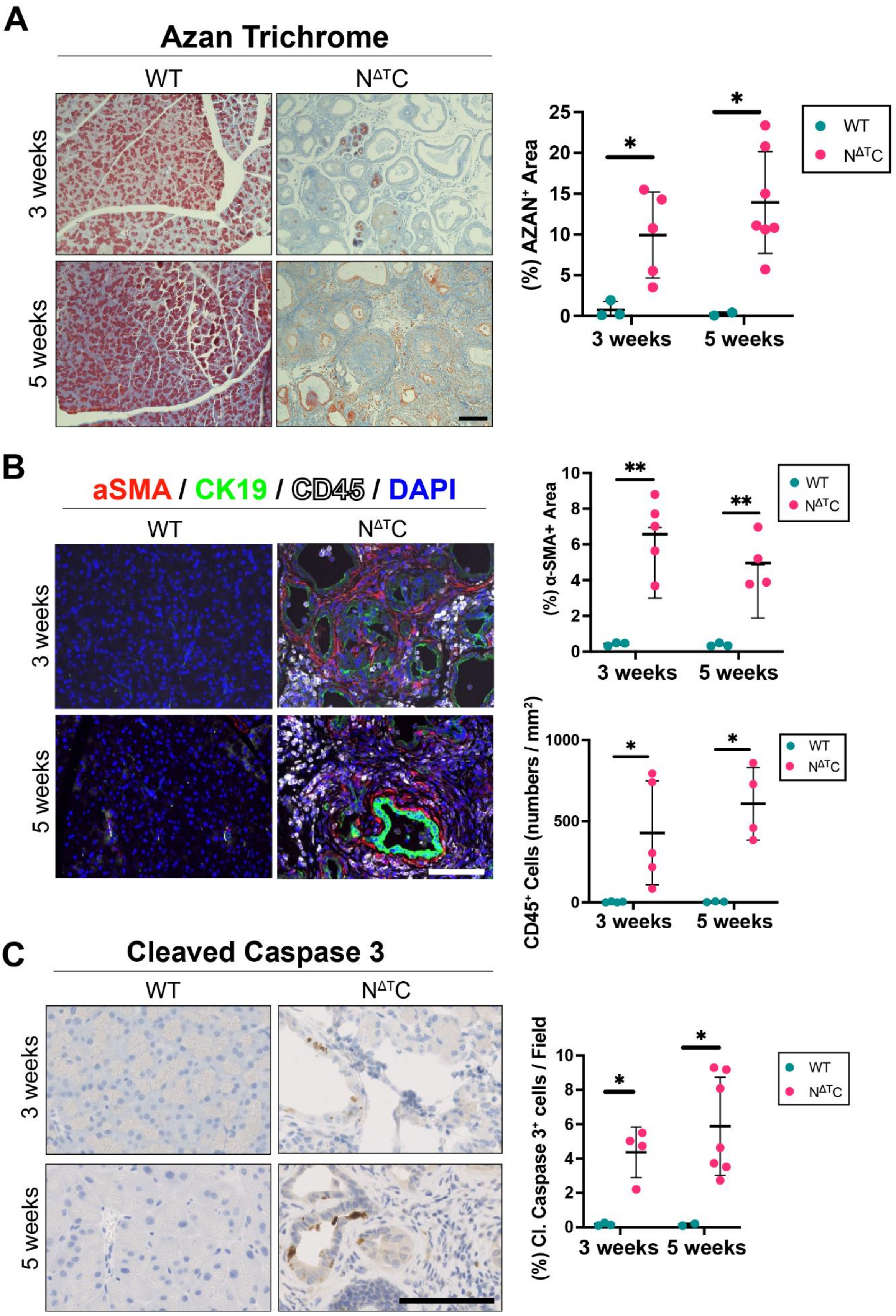


77

78 **Supplementary Figure S3. Stromal activation, immune infiltration, and apoptosis during**
79 **pancreatic remodeling at 3 and 5 weeks.**

**(A) Left:** Representative Azan trichrome staining of pancreatic sections from WT and $N^{\Delta T}C$ mice at 3 and 5 weeks. Scale bars: 100 µm. **Right:** Quantification of fibrotic area as a percentage of the total pancreatic tissue area. N≥2 mice/group. Unpaired t-test. **(B) Left:** Representative immunofluorescence staining of pancreatic sections from WT and $N^{\Delta T}C$ mice at 3 and 5 weeks for α-SMA (red), CK19 (green), CD45 (white), and DAPI (blue). Scale bars: 100 µm. **Upper right:** Quantification of α-SMA-positive area as a percentage of the total pancreatic tissue area. N≥3 mice/group. Welch's t-test. **Lower right:** Quantification of $CD45^+$ immune cells per mm$^2$. N≥3 mice/group. Welch's t-test. **(C) Left:** Representative immunohistochemistry staining for cleaved caspase 3 in pancreatic sections from WT and $N^{\Delta T}C$ mice at 3 and 5 weeks. Scale bars: 100 µm. **Right:** Quantification of cleaved caspase 3-positive cells as a percentage of the total pancreatic tissue area. N≥3 mice/group. Welch's t-test.

Dot plots represent individual mice. Data are presented as mean ± SD. *p < 0.05, **p < 0.01.

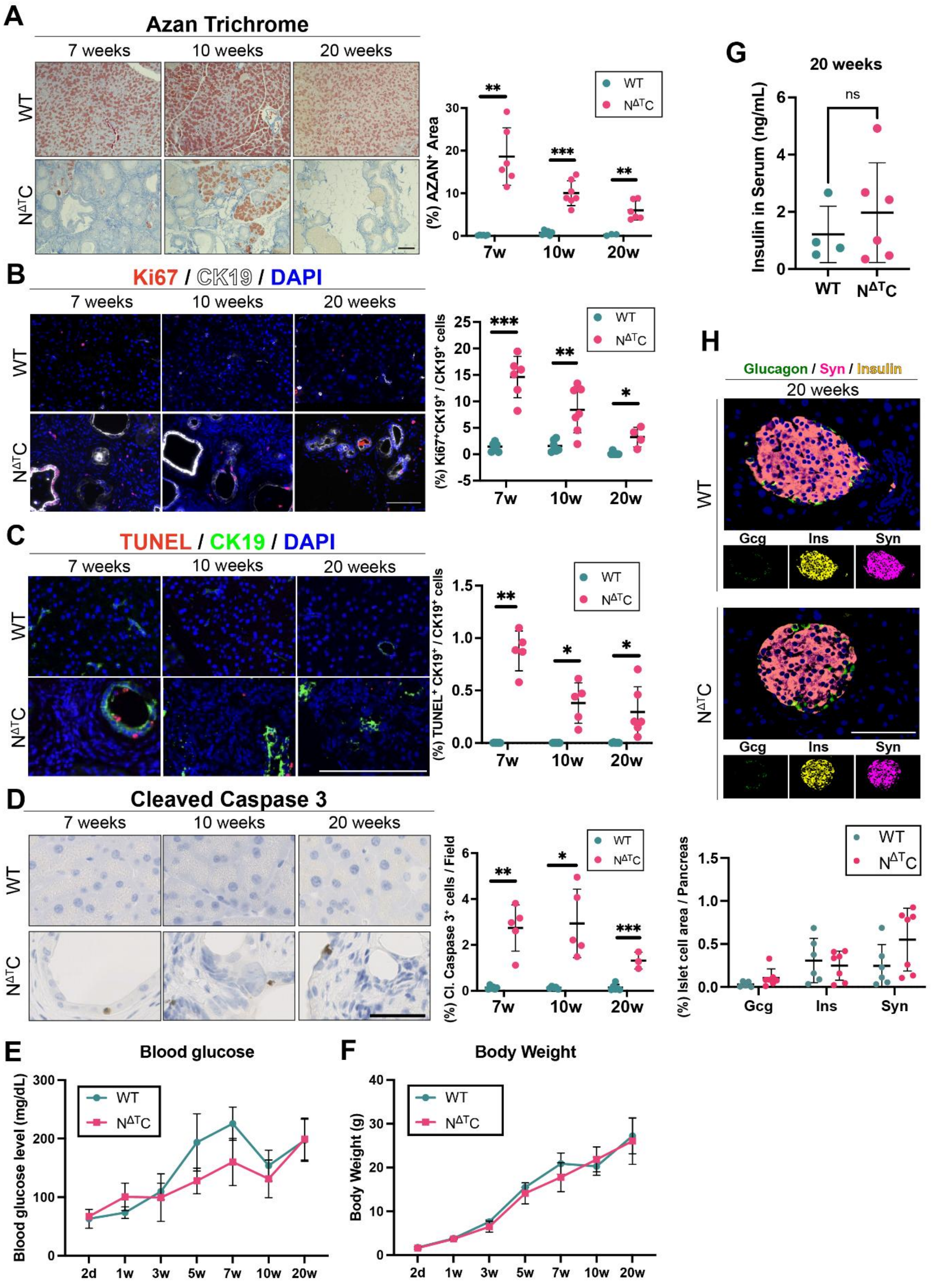


**Supplementary Figure S4**. **Pancreatic remodeling is accompanied by fibrosis, ductal-cell turnover at 7, 10, 20 weeks, and preserved systemic metabolic parameters.**

**(A) Left:** Representative Azan trichrome staining of pancreatic sections from WT and $N^{\Delta T}C$ mice at 7, 10, and 20 weeks. Scale bars: 100 μm. **Right:** Quantification of fibrotic area as a percentage of the total pancreatic tissue area. N≥6 mice/group. Welch's t-test. **(B) Left:** Representative immunofluorescence staining of pancreatic sections from WT and $N^{\Delta T}C$ mice at 7, 10, and 20 weeks for Ki67 (red), CK19 (white), and DAPI (blue). **Right:** Quantification of $Ki67^{+}CK19^{+}$ cells as a percentage of total $CK19^{+}$ cells. N≥4 mice/group. Welch's t-test. Scale bars: 100 μm. **(C) Left:** Representative immunofluorescence staining of pancreatic sections from WT and $N^{\Delta T}C$ mice at 7, 10, and 20 weeks for TUNEL (red), CK19 (green), and DAPI (blue). Scale bars: 100 μm. **Right:** Quantification of $TUNEL^{+}CK19^{+}$ cells as a percentage of total $CK19^{+}$ cells. N≥6 mice/group. 7 weeks: Mann-Whitney test. 10, 20 weeks: Welch's t-test. **(D) Left:** Representative immunohistochemistry staining for cleaved caspase 3 in pancreatic sections from WT and $N^{\Delta T}C$ mice at 7, 10, and 20 weeks. Scale bars: 50 μm. **Right:** Quantification of cleaved caspase 3-positive cells as a percentage of field. N≥3 mice/group. 7 weeks: Mann-Whitney test. 10 weeks: Welch's t-test; 20 weeks: Unpaired t-test. **(E)** Blood glucose levels in WT and $N^{\Delta T}C$ mice at 2 days and 1, 3, 5, 7, 10, and 20 weeks of age. N≥3 mice/group. Upaired t-test. **(F)** Body weights of WT and $N^{\Delta T}C$ mice at 2 days and 1, 3, 5, 7, 10, and 20 weeks of age. N≥3 mice/group. Upaired t-test. **(G)** Serum insulin concentrations in WT and $N^{\Delta T}C$ mice at 20 weeks. N≥3 mice/group. Unpaired t-test. **(H) Top:** Representative immunofluorescence staining of pancreatic sections from WT and $N^{\Delta T}C$ mice for synaptophysin (Syn, red), insulin (Ins, yellow), Glucagon (green) and DAPI (blue). Scale bars: 100 μm. **Bottom:** Quantification of synaptophysin-positive pancreatic area as a percentage of the total pancreatic tissue area. N≥5 mice/group.

Dot plots represent individual mice. Data are presented as mean ± SD. ns: $p > 0.05$, *$p < 0.05$, **$p < 0.01$, ***$p < 0.001$.

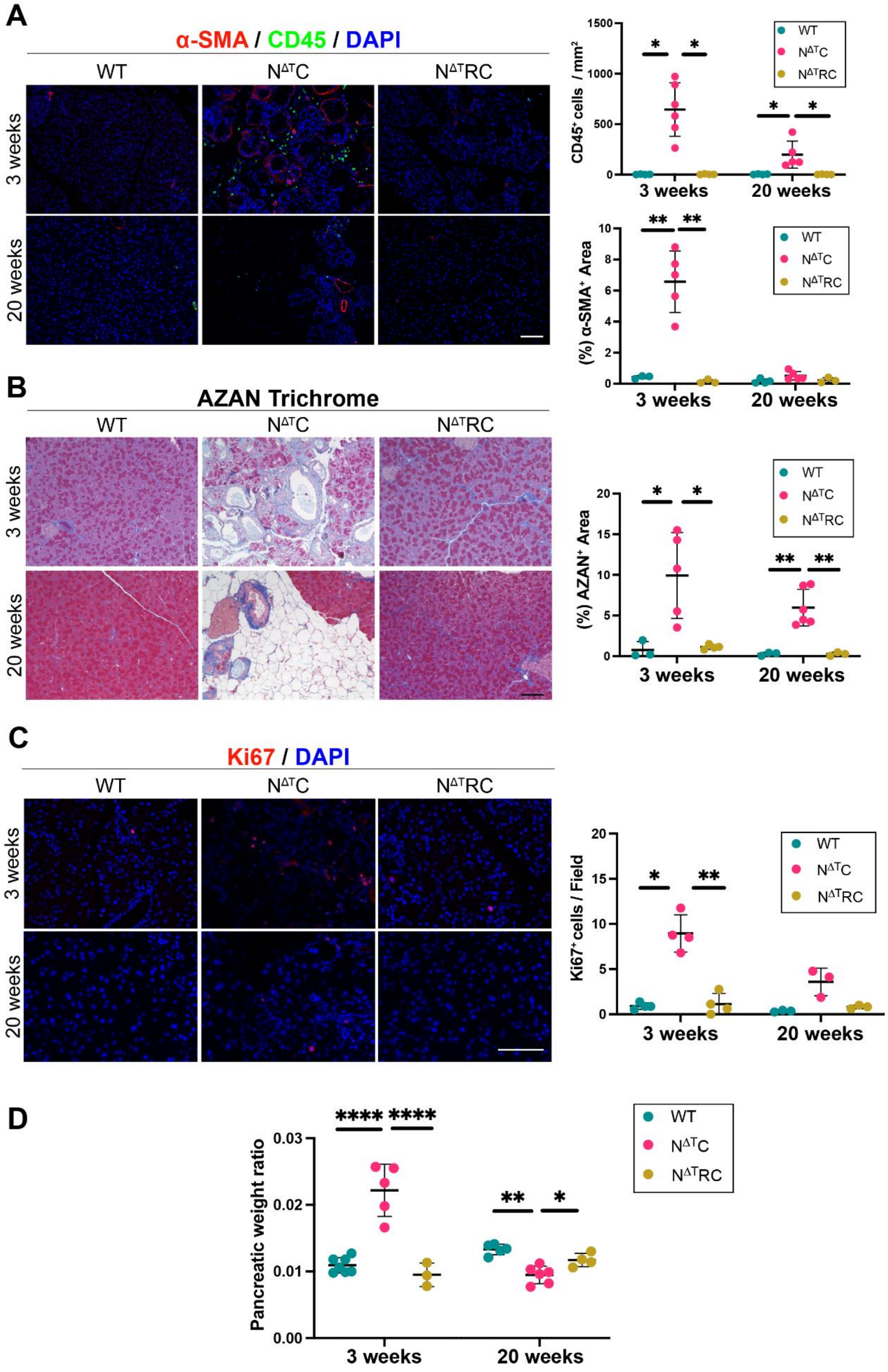


**Supplementary Figure S5. RelB deletion reverses the inflammatory, fibrotic, and proliferative pancreatic phenotype induced by constitutive NIK activation.**

**(A) Left:** Representative immunofluorescence staining of pancreatic sections from WT, $N^{\Delta T}C$, and $N^{\Delta T}RC$ mice at 3 and 20 weeks for α-SMA (red), CD45 (green), and DAPI (blue). **Upper right:** Quantification of $CD45^+$ immune cells per $mm^2$. N≥3 mice/group. 3 weeks: Welch  ANOVA test. 20 weeks: Non-parametric test. **Lower right:** Quantification of α-SMA-positive area as a percentage of the total pancreatic tissue area. N≥3 mice/group. 3 weeks: Welch ANOVA test. 20 weeks: One-way ANOVA test. **(B) Left:** Representative Azan trichrome staining of pancreatic sections from WT, $N^{\Delta T}C$, and $N^{\Delta T}RC$ mice at 3 and 20 weeks. **Right:** Quantification of fibrotic area as a percentage of the total pancreatic tissue area. N≥3 mice/group. Welch ANOVA test. **(C) Left:** Representative immunofluorescence staining of pancreatic sections from WT, $N^{\Delta T}C$, and $N^{\Delta T}RC$ mice at 3 and 20 weeks for Ki67 (red) and DAPI (blue). **Right:** Quantification of $Ki67^+$ cells per field of view. N≥3 mice/group. Welch ANOVA. **(D)** Quantification of the pancreas-to-body weight ratio in WT, $N^{\Delta T}C$, and $N^{\Delta T}RC$ mice at 3 and 20 weeks. N≥3 mice/group**.** One-way ANOVA test.

Dot plots represent individual mice. Data are presented as mean ± SD. *$p < 0.05$, **$p < 0.01$, ****$p < 0.0001$. Scale bars: 100 μm.

**Supplement Table S1:** List of immunofluorescence (IF), immunohistochemistry (IHC) and Western Blot (WB) antibodies.

| Antibody (Dilution) | Catalog Number | Application |
|---|---|---|
| Ki67 (1:200) | Cell Signaling, #9449S | IF |
| aSMA (1:200) | Millipore, #CBL 171 | IF |
| Amylase (1:200) | Sigma, #A8273 | IF |
| CK19 (1:200) | Millipore, #MABT913 | IF |
| CD45 (1:100) | BD Biosciences, #550539 | IF |
| GPA33 (1:100) | Abcam, #ab231592 | IF |
| RFP (1:200) | Invitrogen, #MA5-15257 | IF |
| GFP (1:1000) | Abcam, #ab6658 | IF |
| Glucagon (1:2000) | Novus, #NBP2-21803 | IF |
| Synaptophysin (1:100) | Synaptic Systems, #101006 | IF |
| Insulin (1:100) | Invitrogen, #PA1-26938 | IF |
| Perilipin (1:100) | Abcam, #ab3526 | IF |
| RelB (1:100) | Cell signaling, #10544 | IF |
| Donkey anti-mouse 488 (1:500) | Invitrogen, #A-21202 | IF |
| Donkey anti-mouse 555 (1:500) | Invitrogen, #A-32773 | IF |
| Donkey anti-mouse 594 (1:500) | Invitrogen, #A-21203 | IF |
| Donkey anti-mouse 647 (1:500) | Invitrogen, #A-31573 | IF |
| Donkey anti-rabbit 488 (1:500) | Invitrogen, #A-21206 | IF |
| Donkey anti-rabbit 594 (1:500) | Invitrogen, #A-21207 | IF |
| Donkey anti-rabbit 647 (1:500) | Invitrogen, #A-32795 | IF |
| Donkey anti-rat 488 (1:500) | Invitrogen, #A-21208 | IF |
| Donkey anti-rat 555 (1:500) | Invitrogen, #A-48270 | IF |
| Donkey anti-rat 647 (1:500) | Invitrogen, #A-48272 | IF |
| Donkey anti-goat 488 (1:500) | Invitrogen, #A-11055 | IF |
| Donkey anti-chicken 555 (1:500) | Invitrogen, #A-78949 | IF |
| Goat anti-Guinea pig 647 (1:500) | Invitrogen, #A-21450 | IF |
| Cleaved Caspase 3 (1:100) | Cell Signaling, #9664S | IHC |
| TOM20 (1:1000) | Santa Cruz, #sc-11415 | WB |
| NIK (1:1000) | Santa Cruz, #sc-7211 | WB |
| Anti-rabbit HRP (1:5000) | Santa Cruz, #2004 | WB |
